# Virulence and antibiotic resistance plasticity of *Arcobacter butzleri*: insights on the genomic diversity of an emerging human pathogen

**DOI:** 10.1101/775932

**Authors:** Joana Isidro, Susana Ferreira, Miguel Pinto, Fernanda Domingues, Mónica Oleastro, João Paulo Gomes, Vítor Borges

**Affiliations:** Bioinformatics Unit, National Institute of Health Dr. Ricardo Jorge, Lisbon, Portugal; CICS-UBI-Centro de Investigação em Ciências da Saúde, Universidade da Beira Interior, Covilhã, Portugal; National Reference Laboratory for Gastrointestinal Infections, National Institute of Health Dr. Ricardo Jorge, Lisbon, Portugal

**Author notes:** Corresponding authors: **Susana Ferreira, Email address:**, **Vítor Borges, Email address:**.

**Keywords:** *Arcobacter butzleri*, genome diversity, virulence factors, antibiotic resistance, *porA*, phase variation **Repositories**: Sequence data was deposited in the European Nucleotide Archive (ENA) (BioProject PRJEB34441)

## Abstract

*Arcobacter butzleri* is a food and waterborne bacteria and an emerging human pathogen, frequently displaying a multidrug resistant character. Still, no comprehensive genome-scale comparative analysis has been performed so far, which has limited our knowledge on *A. butzleri* diversification and pathogenicity. Here, we performed a deep genome analysis of *A. butzleri* focused on decoding its core- and pan-genome diversity and specific genetic traits underlying its pathogenic potential and diverse ecology. In total, 49 *A. butzleri* strains (collected from human, animal, food and environmental sources) were screened.

*A. butzleri* (genome size 2.07-2.58 Mbp) revealed a large open pan-genome with 7474 genes (about 50% being singletons) and a small core-genome with 1165 genes. The core-genome is highly diverse (≥55% of the core genes presenting at least 40/49 alleles), being enriched with genes associated with housekeeping functions. In contrast, the accessory genome presented a high proportion of loci with an unknown function, also being particularly overrepresented by genes associated with defence mechanisms. *A. butzleri* revealed a plastic virulome (including newly identified determinants), marked by the differential presence of multiple adaptation-related virulence factors, such as the urease cluster *ureD(AB)CEFG* (phenotypically confirmed), the hypervariable hemagglutinin-encoding *hecA*, a putative type I secretion system (T1SS) harboring another agglutinin potentially related to adherence and a novel VirB/D4 T4SS likely linked to interbacterial competition and cytotoxicity. In addition, *A. butzleri* harbors a large repertoire of efflux pumps (EPs) (ten “core” and nine differentially present) and other antibiotic resistant determinants. We provide the first description of a genetic determinant of macrolides resistance in *A. butzleri*, by associating the inactivation of a TetR repressor (likely regulating an EP) with erythromycin resistance. Fluoroquinolones resistance correlated with the Thr-85-Ile substitution in GyrA and ampicillin resistance was linked to an OXA-15-like β-lactamase. Remarkably, by decoding the polymorphism pattern of the porin- and adhesin-encoding main antigen PorA, this study strongly supports that this pathogen is able to exchange *porA* as a whole and/or hypervariable epitope-encoding regions separately, leading to a multitude of chimeric PorA presentations that can impact pathogen-host interaction during infection. Ultimately, our unprecedented screening of short sequence repeats detected potential phase-variable genes related to adaptation and host/environment interaction, such as lipopolysaccharide modification and motility/chemotaxis, suggesting that phase variation likely modulate *A. butzleri* key adaptive functions.

In summary, this study constitutes a turning point on *A. butzleri* comparative genomics revealing that this human gastrointestinal pathogen is equipped with vast virulence and antibiotic resistance arsenals, which, coupled with its remarkable core- and pan-genome diversity, opens a multitude of phenotypic fingerprints for environmental/host adaptation and pathogenicity.

**IMPACT STATEMENT:** Diarrhoeal diseases are the most common cause of human illness caused by foodborne hazards, but the surveillance of diarrhoeal diseases is biased towards the most commonly searched infectious agents (namely *Campylobacter jejuni* and *C. coli*). In fact, other less studied pathogens are frequently found as the etiological agent when refined non-selective culture conditions are applied. A hallmark example is the diarrhoeal-causing *Arcobacter butzleri* which, despite being also associated with extra-intestinal diseases, such as bacteremia in humans and mastitis in animals, and displaying high rates of antibiotic resistance, has not yet been profoundly investigated regarding its epidemiology, diversity and pathogenicity. To overcome the general lack of knowledge on *A. butzleri* comparative genomics, we provide the first comprehensive genome-scale analysis of *A. butzleri* focused on exploring the intraspecies virulome content and diversity, resistance determinants, as well as how this pathogen shapes its genome towards ecological adaptation and host invasion. The unveiled scenario of *A. butzleri* rampant diversity and plasticity reinforces the pathogenic potential of this food and waterborne hazard, while opening multiple research lines that will certainly contribute to the future development of more robust species-oriented diagnostics and molecular surveillance of *A. butzleri*.

**DATA SUMMARY:** *A. butzleri* raw sequence reads generated in the present study were deposited in the European Nucleotide Archive (ENA) (BioProject PRJEB34441). The assembled contigs (.fasta and .gbk files), the nucleotide sequences of the predicted transcripts (CDS, rRNA, tRNA, tmRNA, misc_RNA) (.ffn files) and the respective amino acid sequences of the translated CDS sequences (.faa files) are available at http://doi.org/10.5281/zenodo.3434222. Detailed ENA accession numbers, as well as the draft genome statistics are described in Table S1.

## INTRODUCTION

*Arcobacter* genus was described by Vandamme et al. (1991) being included in the *Campylobacteraceae* family along with *Campylobacter* and *Sulfurospirillum* genera [1]. The *Arcobacter* genus is vastly distributed over various habitats showing to be a large and heterogeneous group accommodating 29 recognized species [2]. The characterization of new species pointed to the need of a clarification of the genus’ taxonomy, and recently, a genomic comparative analysis of the class *Epsilonproteobacteria* suggested a reclassification of *Arcobacter* genus as a new family *Arcobacteraceae* [3]. Furthermore, a taxonomic and phylogenetic analyses of the 29 recognized species and 11 candidate species led Pérez-Cataluña et al. (2018) [4] to propose a division of the current genus *Arcobacter* in seven different genera: *Arcobacter*, *Aliarcobacter* gen. nov., *Pseudarcobacter* gen. nov., *Halarcobacter* gen. nov., *Malaciobacter* gen. nov., *Poseidonibacter* gen. nov., and Candidate ‘*Arcomarinus’* gen. nov., of which four genera have already be validated [4, 5]. The species *A. butzleri*, *A. cryaerophilus*, *A. skirrowii* and *A. thereius*, all considered human pathogens, will be included in *Aliarcobacter* genus [4]. Amongst these species, the International Commission on Microbiological Specifications for Foods included *A. butzleri* and *A. cryaerophilus* in the list of microbes considered a serious hazard to human health [6]. Moreover*, A. butzleri* has gained attention as an emerging pathogen being associated with enteric diseases, and even bacteremia in humans, and abortion, mastitis or diarrhea in animals [7]. The main symptomatology associated with enteritis in humans caused by *A. butzleri* is persistent diarrhea [8]; however, non-diarrheal illness marked by abdominal pain, nausea, vomiting or fever has been associated with the presence of this bacterium as well [9–11]. In fact, this species has been described as one of the most commonly isolated pathogens in a survey taken in Belgium during the period 2008-2013 where fecal samples from 6774 patients with enteritis were analyzed [10]; other studies rank it as the fourth most common *Campylobacter*-like organism found in human diarrheic samples [8,12–14]. The pathogenic potential of *A. butzleri* still remains to be clarified, nonetheless its relevance is emphasized by the presence of various putative virulence genes, ability to adhere and invade different cell lines, which are important factors for bacterial pathogenicity and establishment of infection [7]. Although disease caused by *A. butzleri* may be self-limited, several case-studies reported the use of antibiotic treatment for intestinal and extra-intestinal infections, mainly from the classes of β-lactams, fluoroquinolones and macrolides [7, 15]. However, the multidrug resistance levels reported may prompt to a possible compromise of the treatment of infections caused by *A. butzleri*, with multidrug resistance rates ranging from 20 to 93.8 % among isolates from animals, food products, environment and human samples [16–21].

The actual role of the putative virulence factors and the mechanisms behind antibiotic resistance are not thoroughly studied, with the lack of information hampering the evaluation of the disease burden of *A. butzleri* and its role as a health hazard.

To date, no comprehensive genome-scale comparative analysis of *A. butzleri* has been performed, which has limited our knowledge on diversification, evolution and pathogenicity of this bacterial species. Here, by performing deep intraspecies comparative genomics of *A. butzleri*, we observed that this emerging pathogen displays a high core-genome diversity and high pan-genome plasticity, leading to multiple virulome/resistome signatures that can markedly shape its pathogenic potential and adaptive capacity.

## METHODS

### *A. butzleri* strains and growth conditions

In order to study the genetic diversity of *A. butzleri*, 49 genomes, including 27 publicly available genomes and 22 newly sequenced isolates from our collection, were selected for the analysis. Our 22 *A. butzleri* strains (Table S1) were isolated from various environments and samples, as slaughterhouse environments and poultry samples [18], meat and vegetables from retail markets [17], raw milks and samples from a dairy plant environment [22] and isolates from river water. The human strain Ab_1426_2003 was isolated in 2003 from a patient with diarrhea (kindly provided by Armélle Ménard from University of Bordeaux). All the strains had been previously identified as *A. butzleri* [17,18,22], with the exception of the aquatic isolates. These were obtained from a 200 mL sample of river water, which was filtered through a 0.22 µm pore diameter membrane and incubated with 9 mL of Arcobacter Broth (Oxoid, Hampshire, England) with CAT Selective Supplement (Oxoid, Hampshire, England). After incubation for 48 h at 30 °C under aerobiosis, 200 µL of enrichment broth was applied onto a 0.45 µm pore diameter membrane filter, which was then placed onto a Blood Agar plate supplemented with 5% (v/v) defibrinated horse blood (Oxoid, Hampshire, England), and allowed to passively filter at room temperature for 30 min. After removal of the filter, plates were incubated at 30 °C for 24 h under aerobiosis. Colonies were selected and identified as described by Vicente-Martins et al. (2018) [17]. Before each assay, *A. butzleri* strains were cultured on blood agar at 30 °C under aerobiosis.

### Antibiotic susceptibility testing

The minimum inhibitory concentration (MIC) of seven antibiotics (ampicillin, cefotaxime, erythromycin, gentamicin, levofloxacin, ciprofloxacin and nalidixic acid) was determined for all strains by the agar dilution method. Briefly, antibiotic dilutions were incorporated in Müeller-Hinton agar supplemented with 5% (V/V) of horse defibrinated blood, and plates were inoculated with 2 µL of each isolate inoculum (turbidity adjusted to 0.5 McFarland and then diluted to ∼10^7^ cfu/mL). After inoculation, plates were incubated for 48 h at 30 °C, under aerobic conditions. Strains were classified as susceptible or resistant according with MIC interpretative criteria for *Campylobacter coli* (erythromycin, gentamycin, ciprofloxacin and nalidixic acid) [23] and for *Enterobacteraceae* (ampicillin, cefotaxime and levofloxacin) [24] (resistance breakpoint: ampicillin >16 µg/mL; cefotaxime >2 µg/mL; erythromycin >8 µg/mL; gentamicin > 2 µg/mL; levofloxacin > 2 µg/mL; ciprofloxacin > 0.5 µg/mL; nalidixic acid >16 µg/mL).

### Urease test

Urease test was performed using a rapid urease test composed of urea broth (Oxoid). The broth was inoculated heavily from a 24 h pure culture and incubated at 37°C for 48 h. A change of color from light orange to pink indicated a positive result.

### Whole-genome sequencing

DNA was extracted on a NucliSens easyMAG platform (bioMérieux), following the Generic 2.0.1 protocol, according to the manufacturer’s instructions. Illumina Nextera XT libraries were subjected to paired-end sequencing (2×150 or 2×250 bp) on an Illumina Miseq platform, according to the manufacturer’s instructions.

### Bioinformatics analysis

#### *A. butzleri* genome selection, assembly and annotation

The list of the 49 genomes selected for this study and respective available metadata is provided in Table S1, as well as the criteria applied for the exclusion of seven out of the 34 genomes available in public repositories (as of June 2018). The 22 newly sequenced genomes and 16 of the 27 publicly available genomes, for which raw reads were available (Table S1), were *de novo* assembled with the INNUca v3.1 pipeline (https://github.com/B-UMMI/INNUca) [25]. Briefly, after reads’ quality analysis (FastQC v0.11.5, http://www.bioinformatics.babraham.ac.uk/projects/fastqc/) and cleaning (Trimmomatic v0.36) [26], genomes were assembled with SPAdes v3.11 [27] and subjected to post-assembly optimization with Pilon v1.18 [28]. *In silico* multilocus sequence typing (MLST) was performed using PubMLST (https://pubmlst.org/arcobacter/) [29]. Due to the heterogeneous depth of coverage observed throughout the genome, all non-redundant contigs excluded according to the INNUca default coverage-based criteria were inspected and kept in the final assembly when samples showed an assembly size loss ≥ 1.8% after INNUca filtering steps, based on the following extra criteria: i) size ≥ 1000 bp; ii) depth of coverage ≥ 1/3 (± 10%) of the whole assembly depth of coverage; iii) confirmation of homology with *A. butzleri* after querying NCBI and/or BIGSI (http://www.bigsi.io/) databases. Prokka v1.12 [30] was applied to annotate the 49 assemblies of the final dataset.

### Global core SNP-based phylogeny

Core genome multi-alignment was performed with Parsnp v1.2 [31] with the parameters *-c* and *-C 2000* and using the genome of strain RM4018 (RefSeq accession number NC_009850) as reference. The core variants alignment generated by Parsnp was used to infer a maximum-likelihood phylogenetic tree using RAxML-NG v0.5.1b [32] with 100 replicates. Gubbins v2.3.1 [33] was used to estimate the impact of recombination on the species phylogeny, using the genome multi-alignment from Parsnp and the best scoring tree from RAxML-NG as input files.

### Gene-by-gene phylogenetic and diversity analysis

Gene-by-gene analysis was performed using chewBBACA v2.0.11 [34], with a training file generated based on the reference genome RM4018 for Prodigal v2.6.3 [35]. Whole-genome MLST (wgMLST) schema (*CreateSchema* module) and allele calling (*AlleleCall* module) were performed using a Blast Score Ratio of 0.6 on all 49 genomes. Only exact and inferred matches were considered, while other allelic classifications (see https://github.com/B-UMMI/chewBBACA/wiki) were assumed as missing loci. After the removal of repeated loci, core-genome MLST (cgMLST) was extracted only considering loci shared by 100% of the genomes (*ExtractCgMLST* module). Both wgMLST and cgMLST data were used to infer pan and core-genome size and assess allelic diversity. This conservative rational increases the reliability of orthologous assignment, although it expectedly led to the mis-exclusion of some genes from core-genome (e.g. hypervariable genes such as *porA* or duplicated genes such as *flaAB*). In order to determine if singletons (i.e. a gene exclusively present in a single genome) were found in genomic clusters, we searched for singletons contiguity, here assumed when two singletons were separated by less than 10500 bp (corresponding to the highest mode length value among the loci in the wgMLST schema). Also, for functional diversity analysis, Clusters of Orthologous Groups of proteins (COGs) categories [36] were assigned to the pan-genome predicted proteins using the script cdd2cog v0.1 [37] after RPS-BLAST+ (Reverse Position-Specific BLAST) (e-value cut-off of 1e-2), where only the best hit (lowest e-value) and first assigned COG were considered. For core loci polymorphism analysis, chewBBACA *SchemaEvaluator* module was used with default parameters to obtain the multiple allele alignments of all core loci. MEGA X [38] was used to calculate the overall mean p-distance (of nucleotide coding sequences, with complete deletion and 1000 replicates) of all core loci alignments, taking advantage of MEGA-CC v10.0.5 [39].

### Identification of virulence and antibiotic resistance determinants

In order to explore the repertoire of genes potentially linked to adaptation and pathogenicity of *A. butzleri*, we constructed a vast database of virulence-, antibiotic resistance- and efflux pump-associated genes (Table S2). The database enrolls: i) virulence determinants already identified by Miller et al [40]; ii) genes identified after querying *A. butzleri* genomes against Resfinder [41], Virulence Factors Database (VFDB) and Comprehensive Antibiotic Resistance Database (CARD) using ABRicate v0.8 (https://github.com/tseemann/abricate); iii) *A. butzleri* homologs of virulence factors found in the closely related human pathogens *Campylobacter jejuni* and *Helicobacter pylori* [42, 43]; iv) and genes potentially associated with efflux pump complex or subunits identified in *A. butzleri* genomes upon chewBBACA-based screening of a CARD sub-database (Antibiotic Resistance Ontology: 3000159). Of note, due to the fact that most of the screened databases contain representative loci from multiple species other than *A. butzleri*, when a hit was identified for a given *A. butzleri* genome, the respective gene/protein homolog of *A. butzleri* was subsequently used to construct the global database to be screened in the remaining genomes. All selected loci were validated with BLASTp against the Non-redundant protein sequences (nr) database to confirm the proteins predicted functionality. The detailed presence/absence and diversity analysis of the selected loci was performed by combining both nucleotide- (using ABRicate) and protein-based (chewBBACA) screening approaches.

### Analysis of the major outer membrane protein PorA

In order to characterize PorA in *A. butzleri*, all *porA* sequences were aligned and the predicted protein sequences were inspected to identify hypervariable regions (HVRs). The sequences of each HVR were subsequently extracted and categorized according to sequence similarity to perform a fine analysis of intra-HVR allelic diversity. Each category enrolls sequences sharing the same length and displaying an overall mean p-distance (regarding amino acid differences) ≤0.2 where each sequence has ≤0.2 p-distance to at least another HVR sequence of the same category. Prediction of the secondary structures of PorA was performed using PRED-TMBB [44] with the posterior decoding method and visualized by TMRPres2D [45], as previously described [46]. BepiPred-2.0 was applied to B-cell epitope prediction [47].

### Identification of phase-variable genes

In order to identify genes potentially regulated by phase-variation mechanisms, *A. butzleri* genomes were screened for the presence of hypermutable simple sequence repeats (SSRs) using PhasomeIt [48]. The cut-off values used were the same as those applied by Aidley et al [48] for *Campylobacter* spp. (-c 7 6 0 5 5 -f W9). Only the SSRs located within or upstream a predicted gene were considered.

## RESULTS

### *A. butzleri* reveals an open pan-genome and a highly diverse core-genome

In the present study, whole-genome sequencing (WGS) was performed for 22 *A. butzleri* strains isolated from multiple sources (human, animal, food and environmental), thus almost doubling the species worldwide genome collection (as of June 2018) (Table S1). The 22 newly sequenced strains show a wide range of resistance profiles. All isolates were resistant to nalidixic acid, 95.4% to cefotaxime (n=21), 45.4% to ampicillin (n=10), 40.9% to levofloxacin and ciprofloxacin (n=9) and 18.1% to erythromycin (n=4); all isolates were susceptible to gentamicin. Overall, nine distinct phenotypic resistance profiles were identified (Table S1).

In total, 49 *A. butzleri* genomes were subjected to deep comparative analysis. *A. butzleri* genome size varied between 2.07 and 2.58 Mbp (with a mean number of 2224 CDSs) and presented a mean GC content of 26.9% (Table S1). The global core SNP-based analysis showed high pairwise variability between all the studied *A. butzleri* isolates, with a mean pairwise distance of 24600 SNPs (ranging from 8187 to 31521 SNPs). The exclusion of putative regions of recombination reduced distances between genomes in ∼30% (mean pairwise distance of 16985 SNPs), but did not lead to significant changes in the inferred phylogenetic tree (Fig. S1). This first insight into *A. butzleri* phylogeny and diversity did not reveal any specific clustering (neither any concordance with geographical location or source), but suggested that the highly diverse studied genomes may illustrate a substantial fraction of the global species diversity. Of note, even though the presence of two copies of the *glyA* locus in *Arcobacter* spp. challenges a proper *in silico* prediction of the MLST profiles with the current short-read sequencing approaches, the strains revealed no shared predicted STs (Table S1).

Gene-by-gene analysis showed that *A. butzleri* harbours an open pan-genome, comprising 7474 loci, with each genome adding a mean of 114 new loci to this large pan-genome scenario. Conversely, the core-genome is small, with 1165 loci (Table S3), and seems to have reached a plateau after adding a few genomes (Fig. 1A). The core-genome presented a very high allelic diversity where more than 55% of the core loci presented ≥ 40 alleles (in a total of 49 strains) (Fig. 1B). Nucleotide polymorphism analysis, however, showed that a high allelic diversity did not always correlate with higher polymorphism (p-distance) (Fig. 1B). Indeed, some loci displaying less allelic diversity (i.e., < 40 alleles) were found among the top polymorphic core loci. For instance, the most polymorphic core locus, *flgD* (ABU_RS09780 homolog; mean pairwise nucleotide p-distance=0.1441), coding for a protein required for flagellar hook formation, presented 35 alleles (Fig. 1B).

**Figure 1.**
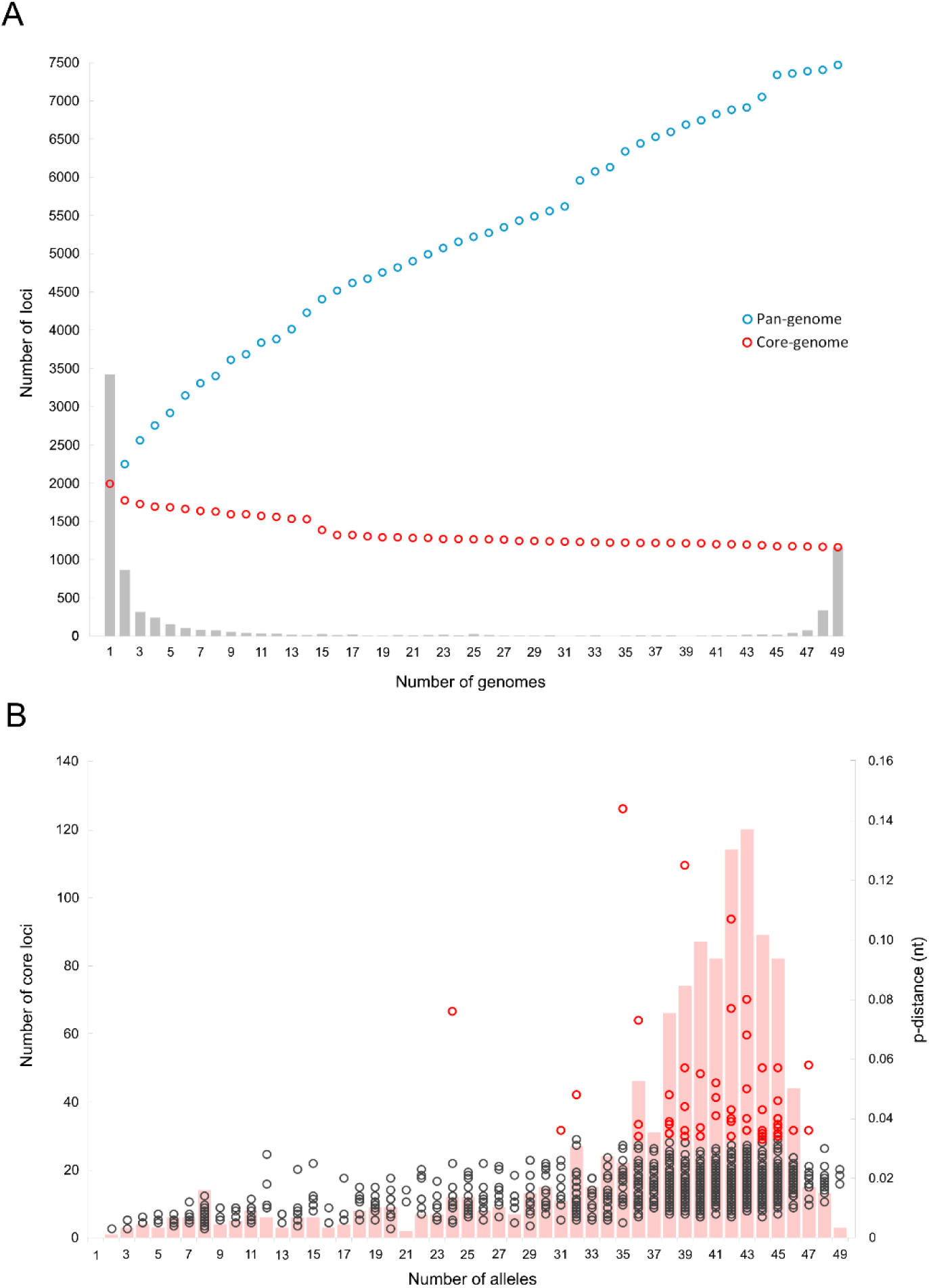
*Arcobacter butzleri* pan- and core-genome. (A) Progression in the number of loci in the pan-genome (blue dots) and core-genome (red dots) as more *A. butzleri* genomes are randomly added to the comparison until a total 49 genomes. Grey bars represent the total number of loci shared by a given number of genomes, with the first bar (Number of genomes = 1) representing the number of singletons and the last bar (Number of genomes = 49) representing the total number of loci shared by all strains, i.e. the core-genome. A representative sequence for each of the 7474 pan-genome loci along with the respective presence/absence profile matrix for all 49 genomes is provided at http://doi.org/10.5281/zenodo.3434222. (B) Relationship between allelic diversity and nucleotide polymorphism of core-genome loci. Pink bars represent the number of loci (left y-axis) harbouring N alleles (x-axis), while dots depict the nucleotide polymorphism (p-distance, right y-axis) detected for each one of the 1165 core loci among the 49 *A. butzleri* strains. The most polymorphic core loci (n=57; refer to methods for details) are marked as red dots. Details, functions and statistics of core-genome loci can be found in Table S3. Nucleotide alignments of the distinct allele sequences detected for each core loci are available at http://doi.org/10.5281/zenodo.3434222.

*A. butzleri* accessory genome comprised 6309 loci, 3433 of which were singletons (i.e. genes exclusively present in a single genome) (Fig. 1A). Overall, singletons accounted for a mean of 3.2% (0.6% - 9.9%) of the genomes total coding capacity. Thirty-two of the 49 genomes contained at least 50 singletons and, among these, three genomes harboured more than 200 singletons (Table S1, Fig. S2). Noteworthy, 90% of all the singletons were found in cluster with at least another singleton, with some genomes presenting remarkably large singleton-enriched clusters. For instance, we found individual clusters in the genomes Ab_CR1132, BMH_AB_252B and Ab_45_11 harbouring a huge proportion of singletons (112, 94 and 74, respectively) which alone accounted for ∼3% of the strains coding capacity. Still, most of these genes were annotated as hypothetical proteins.

### *A. butzleri* core and accessory genomes present distinct functional signatures

In order to inspect the functional diversity of *A. butzleri* pan-genome, all the predicted proteins were classified according to the COG database [36]. Overall, the core and accessory genomes differed greatly in the distribution and proportion of functional categories (Fig. 2). The core genome was enriched with genes associated with housekeeping functions, such as translation or amino acid transport and metabolism (Fig. 2A). Conversely, the accessory genome presented a high proportion of loci with an unknown function (55.4%, contrasting with 21.3% in core-genome) and an expectedly lower number of housekeeping genes (Fig. 2B). This trend was even more pronounced for the strain-specific accessory genome (singletons), where 64.5% of loci had an unknown function. Additionally, it is highlighted that genes related to defense mechanisms are overrepresented among the accessory loci (about 4-fold higher than in core-genome).

**Figure 2.**
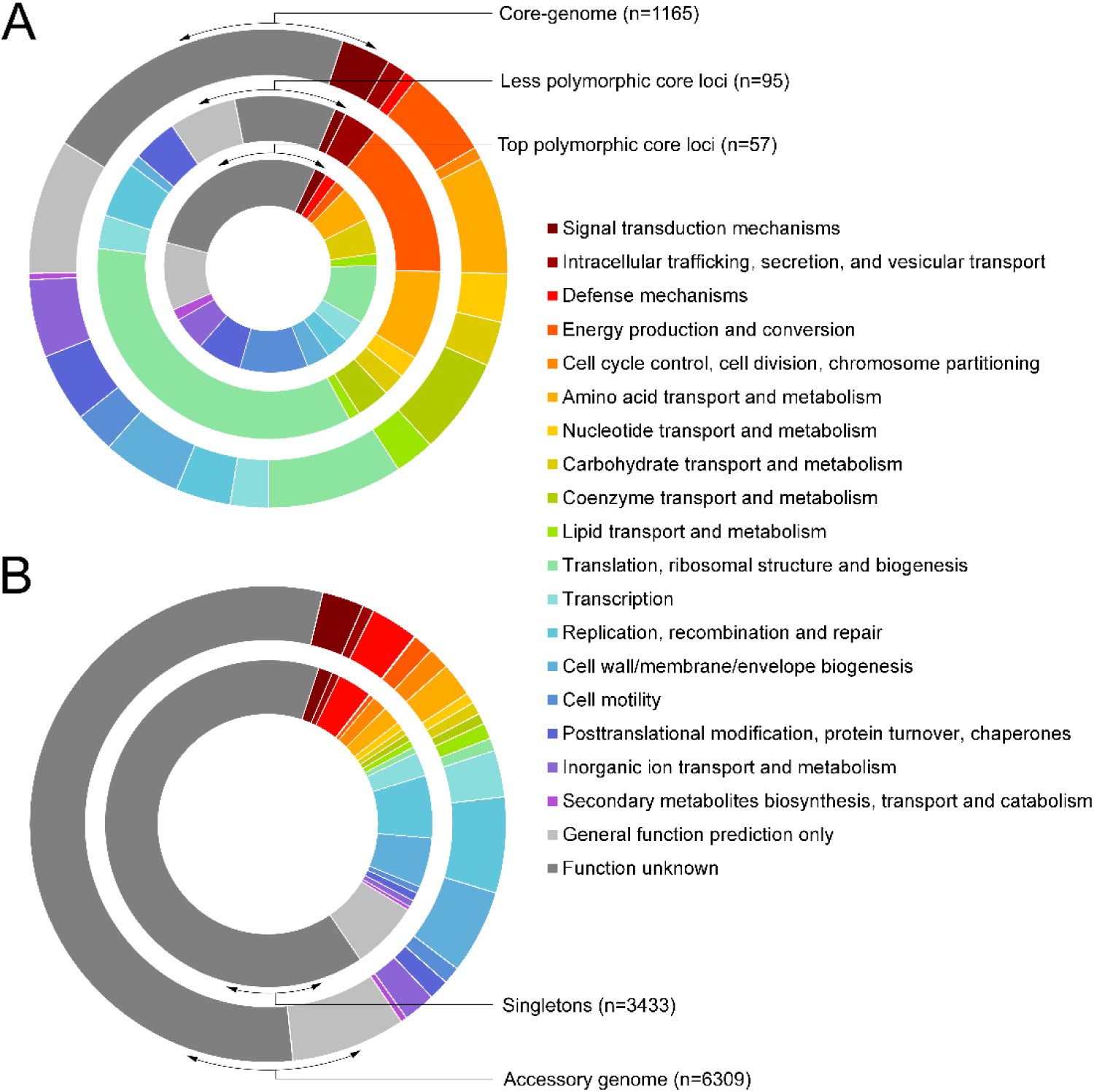
Functional diversity of core and accessory genomes of *Arcobacter butzleri*. (A) Clusters of orthologous groups (COGs) classification of the predicted proteins of the core-genome loci (outer ring), less polymorphic core loci (middle ring) and top polymorphic core loci (inner ring). (B) COGs classification of the predicted proteins of the accessory genome loci (outer ring) and singletons (inner ring).

When correlating the functional category with the degree of polymorphism of the core loci (Fig. 2A), we found that cell motility-associated genes are overrepresented among the top polymorphic core loci when compared to all core loci (10.5% versus 2.8%, respectively). Conversely, most of the less polymorphic core loci (<0.5-fold the median p-distance of all core loci) were associated with translation, ribosomal structure and biogenesis (34.7%) and energy production and conversion (14.7%).

### *A. butzleri* presents a vast arsenal of potential virulence factors

The emerging human pathogen *A. butzleri* is a member of the Epsilonproteobacteria along with some pathogenically relevant and well-characterized species, like *C. jejuni* and *H.pylori*. In this context, *A. butzleri* genomes were queried against a custom database of genes potentially linked to adaptation and pathogenicity (Table S2). *A. butzleri* is a motile bacterium that exhibits a polar flagellum, which is a well-known virulence trait [49]. Here, we observed that most flagellar genes are clustered in two main genomic regions enrolling 20 (ABU_RS09675-ABU_RS09825) and eight (ABU_RS01005-ABU_RS01060) genes. Six additional flagellar genes were found outside these regions, namely, *flaA* and *flaB*, encoding the major and minor flagellin subunits, respectively (ABU_RS11245, ABU_RS11250), *motA* and *motB* (ABU_RS01995, ABU_RS02000), *fliP* (ABU_RS04985) and another flagellin coding gene, *hag* (ABU_RS05570). These 34 flagellar genes were present in all the genomes analyzed; however, it is noteworthy that the genes *fliM* (ABU_RS01005; encoding a flagellar motor switch protein) and *flgE1* (ABU_RS09770; encoding a flagellar hook protein) revealed truncating mutations in strains Ab_1426_2003 and Ab_L352, respectively, potentially impacting the flagellar assembly/function. Remarkably, six of the flagellar genes were found among the top polymorphic core genes, namely *flgD*, *flgL*, *flgK*, *flgE2*, *flgG2* and *flgH* (Table S3).

Among other potential virulence factors found to be present in all the studied genomes (Table S2), we highlight: i) chemotaxis system genes *cheA-cheY* (including three *cheY* genes) [40], with the genome BMH_AB_245B harbouring a frameshift mutation, that leads to an early stop codon in ABU_RS05920 (*cheY*); ii) several genes potentially associated with cellular adherence and invasion, namely *ciaB* (host cell invasion in *Campylobacter* spp.), *tlyA* (hemolysin with a role in cellular adherence in *C. jejuni*), *cadF* and *cj1349* (fibronectin binding proteins, that promote bacteria to cell contact) and *pldA* (phospholipase associated with lysis of erythrocytes) [40, 50]; iii) and several homologs of *Campylobacter* spp. virulence factors [42], namely *luxS* (chemotaxis-associated), *htrA* (chaperone involved in adhesins folding); *iamA* (invasion-associated gene), *fur* (ferric uptake regulator), and *mviN* (inner membrane protein required for peptidoglycan biosynthesis).

Besides this repertoire of “core” virulence factors, we additionally found multiple genes potentially linked to pathogenicity and adaptation that were differentially present among *A. butzleri* strains (Fig. 3, Table S2). We found the chemotaxis-associated *docA* homolog [51] in six *A. butzleri* strains. Of note, the *docA* adjacent gene in *C. jejuni*, *docB*, also a potential determinant of chick colonization [52], was not found among the studied strains. The absence of *docA* in most strains seems to find a parallel in *C. coli* (which also usually lacks *docB*) but not in *C. jejuni*, that usually carries both genes [51]. The putative virulence determinants *irgA* and *iroE*, which may be linked to uropathogenicity in *Escherichia coli* [40,53–55], were both found (contiguously) in 21 strains, with *irgA* being present alone in eight additional genomes. The *Campylobacter* spp*. cfrB* homolog, involved in iron uptake [42], was identified in 18 genomes. Noteworthy, *hecA* and *hecB*, which code for a filamentous hemagglutinin and a hemolysin activation protein, respectively, were always found simultaneously (and adjacent) in a total of 13 genomes. Contrarily to *hecB*, *hecA* was found to be extremely polymorphic (yielding hypervariable predicted proteins, both in sequence and size) which challenges *in silico* WGS-based screening strategies. Indeed, exclusive direct nucleotide homology screening would yield false negatives for *hecA* presence (Fig. 3). This is also relevant since *hecB* is more frequently found than *hecA* in rapid PCR-based screening of *A. butzleri* potential virulence [21,56–60]. Concordantly, we observed a low homology between the commonly used primers [56] and most of *hecA* sequences found in the studied genomes. Furthermore, resembling *H. pylori*, *A. butzleri* contains a urease cluster, enrolling six genes, *ureD(AB)CEFG*, which was detected in 25 genomes. Our 22 *A. butzleri* isolates (10 carrying the urease cluster) were subjected to a functional urease assay that confirmed the urease-positive phenotype in 9 out of the 10 strains containing the urease cluster. A fine sequence inspection of the urease cluster of the strain phenotypically testing negative (Ab_CR891) revealed several exclusive genetic traits that may justify the observed incongruence: i) an alternative start codon (Val instead of Met) in the *ureAB*; ii) multiple protein-changing mutations in *ureC*, which encodes the alpha urease subunit, with most of them occurring in close proximity with residues involved in subunit interactions (Fig. S3). Also of note, a cluster of seven genes potentially associated with capsule polysaccharide biosynthesis, including a homolog of the *C. jejuni gmhA2*, was identified in a single strain (Ab_4511). Finally, we highlight the detection, in a single strain (D4963), of a complete type IV secretion system (T4SS), resembling a VirB/D4 secretion system (i.e., *virB1-11* and *virD4* genes, including the pathogenicity island-associated *cag12* homolog, *virB7*), which carries a large putative adhesin-encoding gene (D4963p_10810) and toxin/anti-toxin systems (Fig. 3, Table S2). This 56 kb cluster comprises 56 genes (D4963p_10540 - D4963p_11080) and is inserted in a D4963 genome region where the reference strain RM4018 harbors a distinct ∼16 kb mobile element (between the ABU_RS08535 and ABU_RS08605 loci).

**Figure 3.**
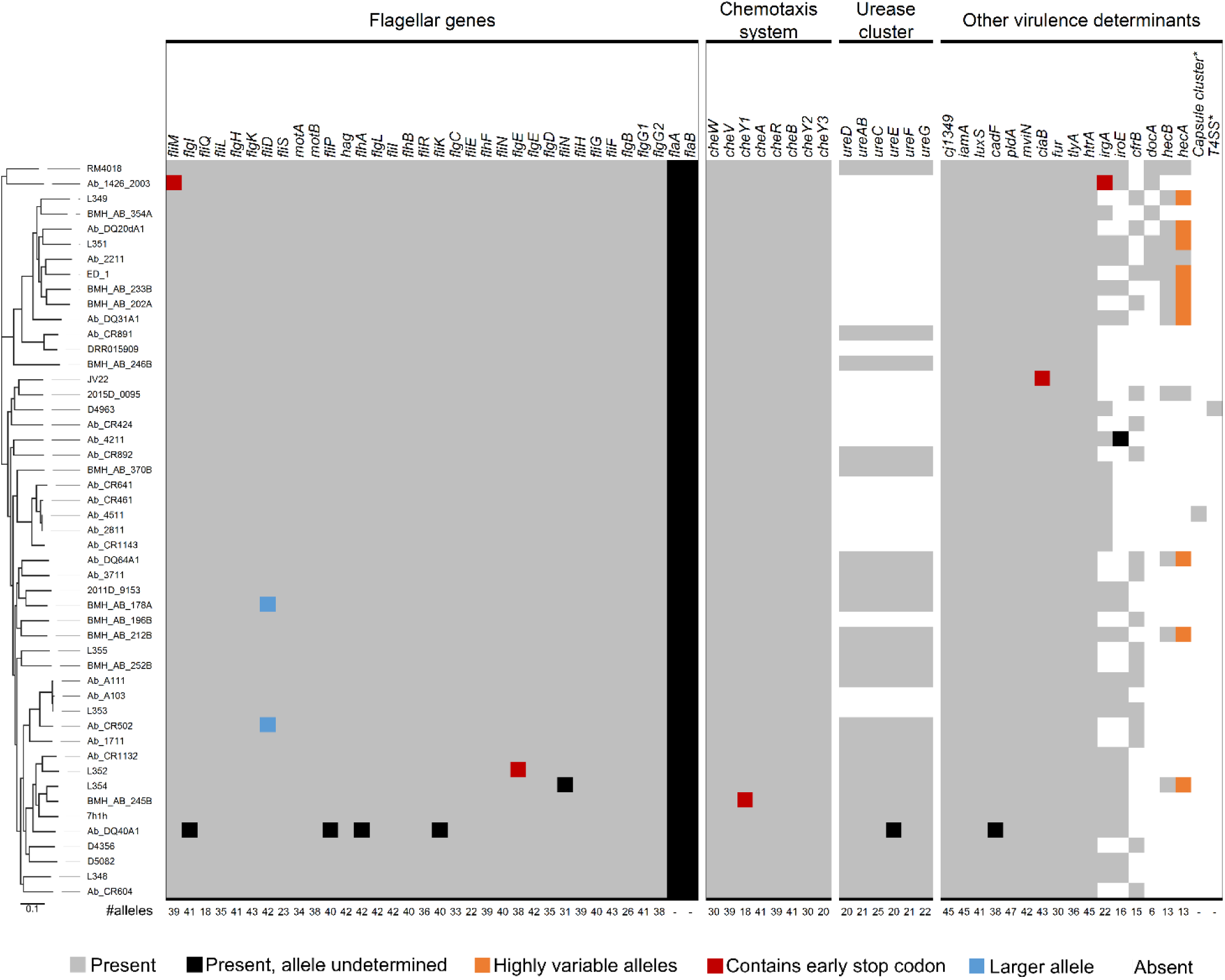
The vast arsenal of *Arcobacter butzleri* virulence factors. Presence/absence of multiple virulence factors identified among the 49 genomes studied (organized according to their position in the core-genome SNP-based phylogeny after removal of predicted recombinant regions), including 34 flagellar genes (some of which associated with the type III secretion system), chemotaxis-associated genes, the urease encoding cluster *ureD(AB)CEFG*, and other virulence determinants. The numbers placed under each column represent the total number of alleles identified for each locus (not determined for the duplicated *flaA* and *flaB* genes). *The putative capsule-related cluster (only present in strain Ab_45_11) and the type IV secretion system (T4SS)-containing region (only present in strain D4963) enroll seven and 56 genes, respectively, and are presented as single blocks for simplicity. Refer to Table S2 for details on the depicted loci and to Figure 5 for details on the genotype-phenotype associations established in this study.

### *A. butzleri* harbors a large repertoire of efflux pump-related genes and other antibiotic resistant determinants

Nineteen putative efflux pump (EP) systems (hereinafter designated EP1-EP19) belonging to different families and enrolling a total of 56 genes could be detected in the 49 *A. butzleri* genomes (Fig. 4, Table S2). Ten of these transporters were present in all the genomes: i) EP2, EP12 (located immediately downstream of *flhA*), EP13 and EP14, belonging to the major facilitator superfamily (MFS); ii) EP8, a transporter from the small multidrug resistance (SMR) family; iii) EP5, EP6 and EP10, belonging to the ATP-binding cassette (ABC) superfamily, with EP10 sharing homology with the YbhGFSR transporter previously described in *E. coli* [61] and EP5 corresponding to a lipid A export ATP-binding/permease protein MsbA (encoded by a top polymorphic core gene); and iv) EP4 and EP15, members of the resistance-nodulation-division (RND) family. Of note, EP4 is located downstream of the urease cluster in the genomes containing this operon.

**Figure 4.**
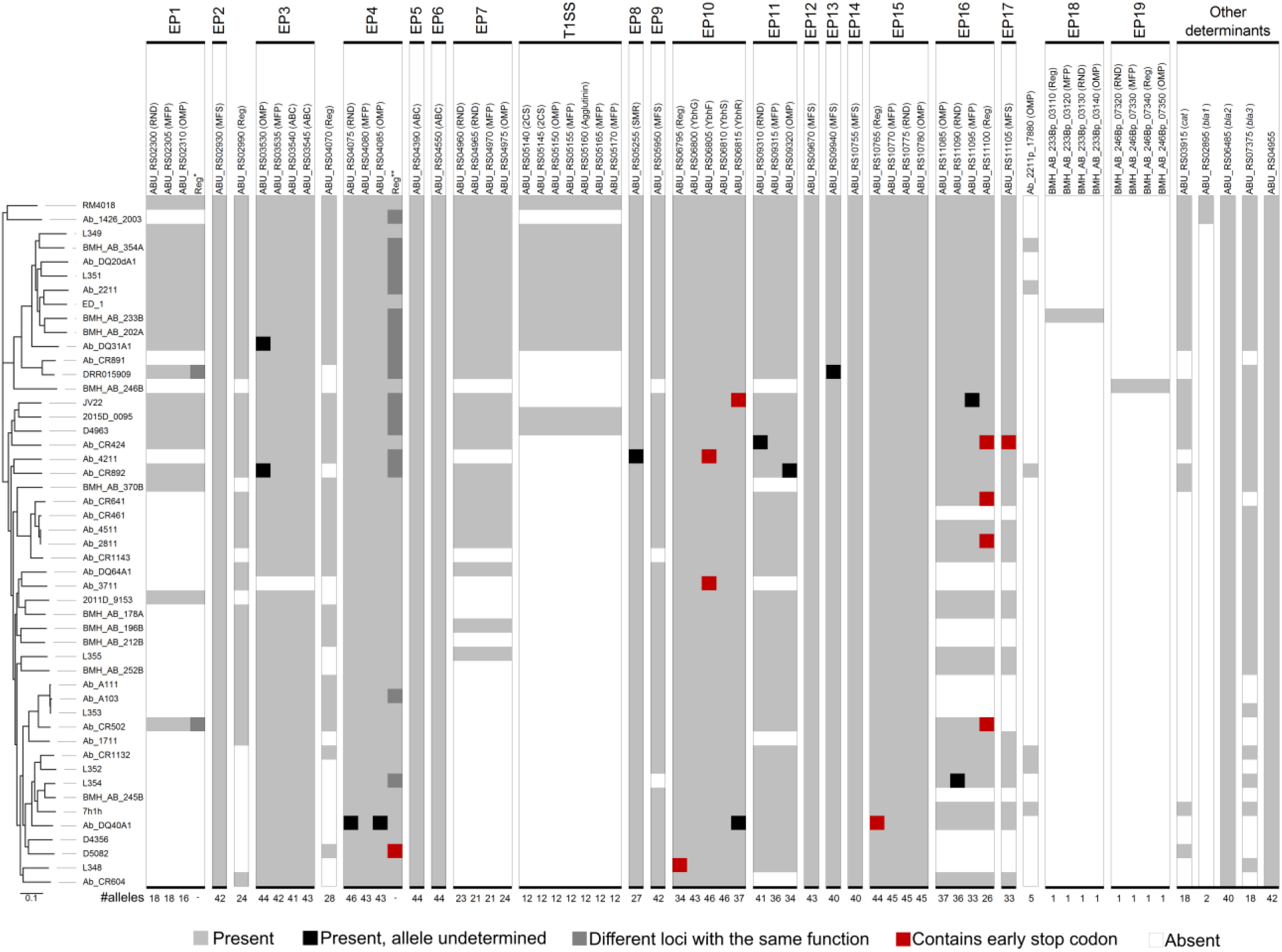
The large repertoire of efflux-pump-related genes and other antibiotic resistant determinants identified in *Arcobacter butzleri* genomes. Presence/absence of multiple efflux-pump-related genes identified among the 49 genomes studied (organized according to their position in the core-genome SNP-based phylogeny after removal of recombination), including 19 efflux-pumps (EP1-EP19), enrolling one to five genes each, and a putative type I secretion system (T1SS), shown according to their order in the reference genome of strain RM4018. The predicted function/homology is given for each gene, next to the locus tag. The truncating mutations in the TetR regulator of EP16 indicated in four strains are associated with erythromycin resistance. Other resistance determinants are also shown, including three β-lactamase encoding genes (*bla1-bla3*), one of which is the OXA-15 β-lactamase associated with ampicillin resistance (*bla3*), and a chloramphenicol acetyltransferase. *For two strains, a different regulatory gene is present and corresponds to the loci Ab_CR502p_11350 and DRR015909p_21790. **Some strains contain an alternative loci as regulator in this EP (homolog of the gene Ab_A103p_18290). Reg – regulator, MFP – membrane fusion protein, RND – resistance-nodulation-division subunit, OMP – outer membrane protein, ABC - ATP-binding cassette, MFS – major facilitator superfamily, SMR - small multidrug resistance family. Refer to Table S2 for details on the depicted loci and to Figure 5 for details on the genotype-phenotype associations established in this study.

Among the nine EP systems that are not shared by all the 49 *A. butzleri* strains (Fig. 4, Table S2), only two (EP18 and EP19, both RND efflux pumps) were found to be exclusive of a single strain. The remaining seven EP systems belong to the RND, ABC and MFS families, each being present in at least 19 genomes (Fig. 4). We highlight EP16, since the protein size of its regulator TetR (ABU_RS11100) could be correlated with the erythromycin resistance phenotype (Fig. 4, Fig. 5, Table S1). In fact, the four erythromycin resistant strains revealed truncated predicted proteins with 116-142-aa (instead of the 179-aa full length protein) due to a SNP, small indels or an insertion sequence (IS) element (Supplementary Dataset X). In fact, for one strain (Ab_CR424), the inactivating event involved the introduction of a not previously described 2632-bp IS element (harbouring two putative transposases, Ab_CR424p_22200 and Ab_CR424p_22210) in the 352-353-bp position of the TetR-coding gene (manually assembled *tetR* plus IS element can be found in Supplementary Dataset X). The IS revealed imperfect inverted repeats in each end and shares a small direct repeat with the insertion site in *tetR*. Besides these EP systems, we highlight the presence of a potential T1SS enrolling not only HlyD-like and regulatory proteins, but also the multidrug efflux associated protein TolC (not usually found in the Hly operon [62]) and a large protein (2517-aa) resembling a secreted agglutinin RTX potentially related to invasion and adhesion functions [63]. Remarkably, the 12 strains carrying this T1SS are the only ones displaying all the EP-systems detected in the present study (with the exception of the two singleton EP-systems). Nine out of these 12 strains segregate together in the core-genome phylogeny still they show a considerable core-genome background diversity and no metadata is available supporting a link between these strains (Fig. 4).

**Figure 5.**
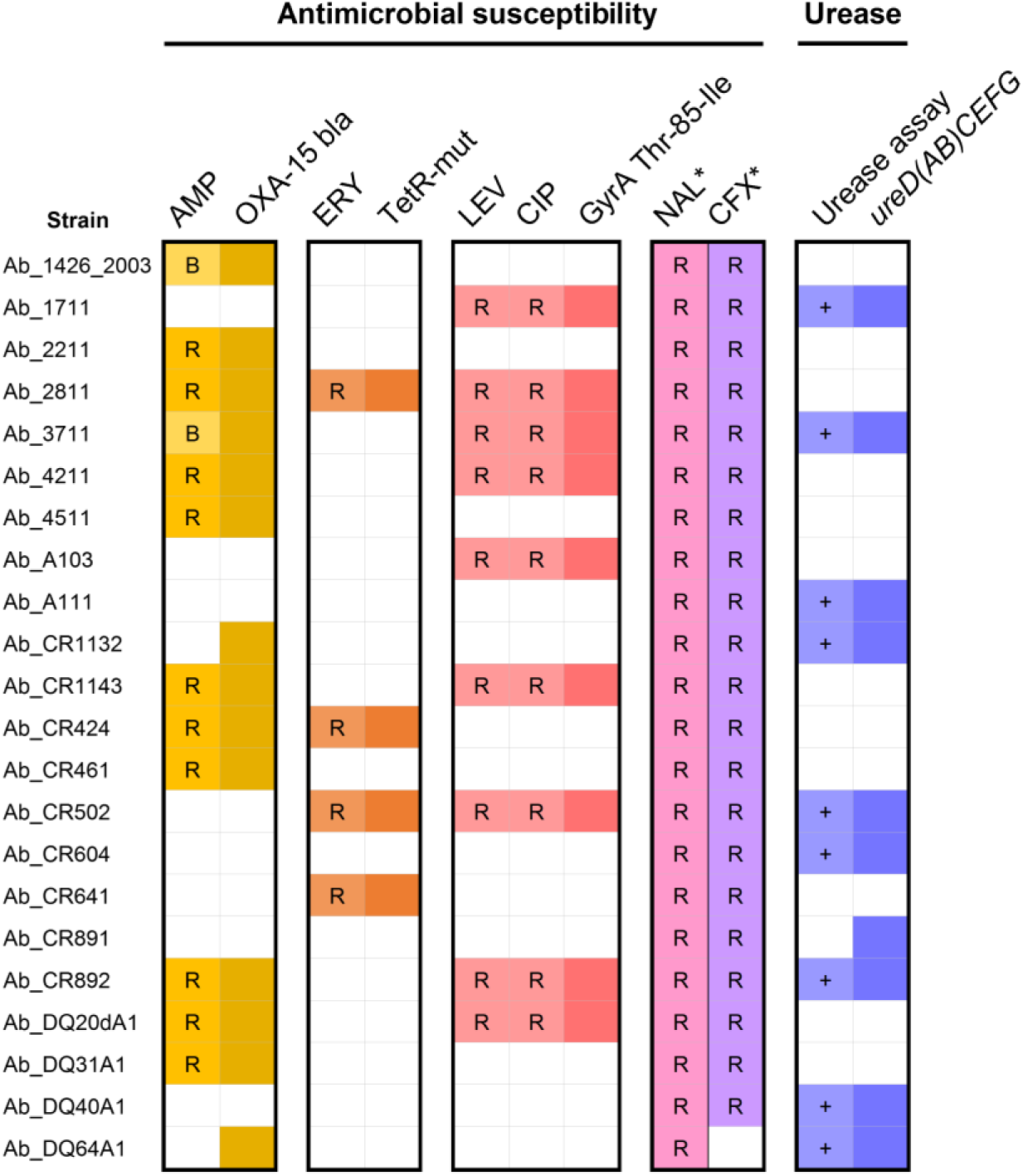
Antibiotic susceptibility and genotype-phenotype associations established in this study. For the 22 phenotypically characterized strains from our collection, each phenotypic trait (antibiotic susceptibility and urease assay) is presented adjacently to its associated genotype, whose presence/absence is indicated by colored/white blocks. *For nalidixic acid and cefotaxime no genotype-phenotype association could be determined. R - resistant phenotype, B - borderline resistant phenotype, + - positive urease assay, AMP - ampicillin, ERY - erythromycin, LEV - levofloxacin, CIP - ciprofloxacin, NAL - nalidixic acid, CFX - cefotaxime.

Finally, the survey of additional antibiotic resistance determinants for the strains of our collection (n=22) revealed the presence of the Thr-85-Ile substitution in the quinolone-resistance determining region of GyrA in all the fluoroquinolones (levofloxacin and ciprofloxacin) resistant strains, as well as a strong correlation between reduced susceptibility to ampicillin and the presence of an OXA-15-like β-lactamase (ABU_RS07375, *bla3*) (among the 14 strains harboring the gene, 10 are resistant and two revealed borderline MICs) (Fig. 4, Fig. 5). In the absence of a strain susceptible to nalidixic acid, no genetic determinants could be specifically associated with resistance to this quinolone. For cefotaxime, the only susceptible strain (Ab_DQ64A1) shares the same presence/absence profile of β-lactamases as other resistant strains and presents multiple strain-specific genetic features. This, together with the absence of phenotypic data for the other publicly available genomes (Table S1), hampered the establishment of a genotype-phenotype association for these antibiotics. Of note, our screening of other potential genetic determinants also detected genes encoding two additional β-lactamases (ABU_RS02895 and ABU_RS06485) and a chloramphenicol acetyltransferase (*cat*, ABU_RS03915) (Fig. 4).

### PorA reveals a rampant mosaicism mediating multiple antigenic fingerprints

The major outer membrane protein and main antigen PorA is known to play a key role in the virulence and adaptation of multiple bacterial pathogens, including *C. jejuni*. As such, we sought to evaluate the genetic diversity of *porA* among *A. butzleri* strains (ABU_RS10145). Not unexpectedly, PorA revealed hypervariable regions (HVRs), both in sequence content and size, alternating with conserved regions. Positional mapping of HVRs against the *C. jejuni* PorA [46] corroborates HVRs localization in externally exposed epitope-containing loops (Fig. 6A). The prediction of transmembrane strands and epitope-enriched regions in *A. butzleri* PorA further supports that HVRs encode externally exposed antigenic regions (Fig. 6A, Fig. S4). HVR3, corresponding to the *C. jejuni* highly virulence-associated external loop 4 [46], revealed the greatest diversity with a total of 33 distinct sequences ranging from 10 to 59-aa. Remarkably, although we found 44 non-redundant PorA protein sequences, identical or highly similar *porA* sequences were detected in strains that are highly distant at phylogenetic level, sustaining the occurrence of homologous recombination of *porA* (Fig. S5A). Likewise, genetic exchange likely mediated the introduction of highly distinct *porA* allelic profiles in closely related strains (Fig. S5B). Homologous recombination frequently involved *porA* and flanking regions, in particular two downstream loci coding for a toxin/anti-toxin system (Fig. S5A). Subsequently, we sought to inspect if individual HVRs were also exchanged between strains. For this, the amino acid sequences of each HVR were categorized by similarity, with each category including same-length sequences with at least 80% pairwise homology (see details in methods). We observed groups of strains carrying highly similar PorA with the exception of discrete HVRs, sustaining an additional scenario of rampant interstrain exchange of specific (or combination of) HVRs (Fig. 6B, Fig. S5C). Altogether, our results sustain a scenario of *porA* diversification marked by ancestral recombination of the whole gene followed by (or simultaneously with) genetic exchange of discrete regions between strains sharing a more similar *porA* conserved backbone, regardless of their genome relatedness (Fig. 6B).

**Figure 6.**
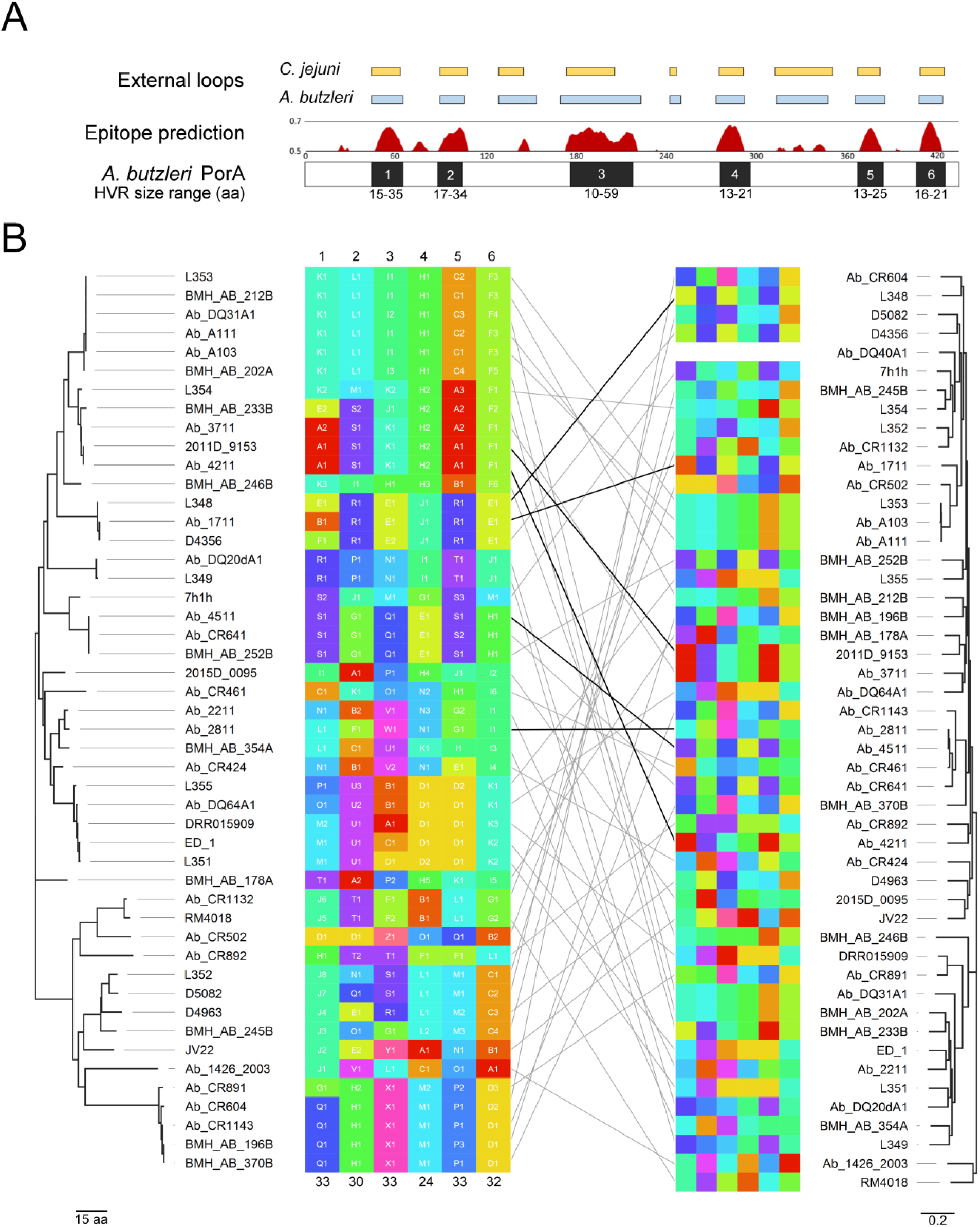
The puzzling character of *Arcobacter butzleri* main antigen PorA. (A) Prediction of the localization of the external loops of *A. butzleri* PorA (blue), with comparison to *Campylobacter jejuni* PorA (yellow), and prediction of B-cell epitope enriched regions (only residues with ≥0.5 predicted probability of being epitope-associated are shown). Residue position is indicated in the x-axis and is relative to the reference strain RM4018 PorA. The relative localization of *A. butzleri* PorA hypervariable regions HVR1-6 is shown in black with the amino acid size range of each HVR being indicated below the corresponding HVR. (B) *A. butzleri* PorA HVRs were categorized according to size and sequence similarity. For each HVR (colored columns 1-6), each category enrolls sequences (amino acid) sharing the same length and displaying an overall mean p-distance ≤ 0.2 and are displayed with the same color and letter. Within each category, different numbers identify distinct amino acid sequences. These HVR mosaics are organized based on the strains relatedness according to: *left panel*) the inferred phylogeny of the non-hypervariable regions of PorA (based on 145 amino acid variant sites); *right panel*) the core-genome SNP-based phylogeny (based on 114777 variant sites detected after recombination removal). Cross-panel lines link the same strain in both panels, where black lines highlight the genomes selected for fine-tuned analysis of recombination (details in Figure S5). The numbers below the left colored columns indicated the number of non-redundant amino acid sequences detected for each HVR. Note: The PorA of strain Ab_DQ40A1 was excluded from this analysis since *porA* was misassembled.

### Phase variation mechanisms likely modulate *A. butzleri* key adaptive functions

Variability of simple sequence repeats (SSRs) is known to mediate important phase-variable phenotypic changes with impact on adaptation capability and pathogenicity in several bacteria, including *C. jejuni* and *H. pylori* [64]. Here, among the 49 *A. butzleri* genomes, we detected 138 genes associated with SSRs (one SSR per gene, with the exception of three genes linked to more than one SSR type) (Table S4). The detected SSRs are located either within or upstream the coding region in similar proportions and include 106 homopolymeric tracts (65 polyA/T and 41 polyG/C), 24 2-bp tandem repeats and 13 4-5-bp tandem repeats. Of note, most of the SSRs were found in a single genome, although the potentially affected genes could be present in multiple genomes but without any associated SSR. A functional assignment of all SSR-related genes revealed an overrepresentation of functional categories (COGs) commonly linked to phase-variation, namely cell wall/membrane/envelope biogenesis, lipid transport and metabolism and inorganic ion transport and metabolism. Still, most genes code for proteins with an unknown function (Table S4). In order to identify SSRs with a higher likelihood of mediating phase-variation mechanisms, each SSR (and the corresponding gene) present in more than one genome were inspected for the presence of interstrain length/sequence variation. Notably, we identified six genes with interstrain in-length variable SSRs (five of which were polyG/C) located within the coding region and potentially leading to ON-OFF switching of protein status (Table 1). We also found three genes presenting either in-length variable SSRs and/or different types of SSRs in the upstream region that may affect gene expression. Noteworthy, the proteins coded by this set of potential phase-variable genes enrol bacterial functions related to adaptation and host/environment interaction, such as lipopolysaccharide (LPS) modification and motility/chemotaxis (Table 1).

**Table 1.** Genes presenting simple sequence repeats with interstrain in-length variability.

| Representative loci | Predicted function | Location of SSR | N genomes with SSR | SSR Tract (number of repeats) | Evidence of ON/OFF status |
| --- | --- | --- | --- | --- | --- |
| Ab_1711p_14260 | Peptidoglycan/LPS O-acetylase | Within ORF | 2 | G(7,8) | ON - 669 aa (G7) ; OFF - 467 aa (G8) |
| Ab_3711p_17430 | Peptidoglycan/LPS O-acetylase | Within ORF | 2 | G(7,9) | ON - 330 aa*; OFF - 166, 171 aa (G7,9) |
| Ab_CR641p_09510 | Peptidoglycan/LPS O-acetylase | Within ORF | 2 | G(7,8) | - |
| Ab_CR604p_16570 | Methyltransferase | Within ORF | 2 | G(7,8) | ON - 283 aa (G7); OFF - 164 aa (G8) |
| Ab_CR1132p_13010 | N-acetyltransferase | Within ORF | 2 | G(7,9) | - |
| Ab_DQ20dA1p_22180 | Hypothetical protein | Within ORF | 3 | AT(7,9) | ON - 107 aa (AT7); OFF - 68 aa (AT9) |
| Ab_1711p_21780 | Response regulator | Upstream | 8 | TA(5,6) | NA |
| Ab_A103p_12080 | Methyl-accepting chemotaxis protein | Upstream | 3 | AAGAT(1,2,5), TAAAA(5,8,21) | NA |
| Ab_CR502p_15020 | Hypothetical protein | Upstream | 6 | ATAAA(1,6,8,12,22), AAGAT(1,2,5) | NA |

SSR – simple sequence repeat, aa – amino acid, NA - not applicable, \*ON protein length corresponds to homolog gene without SSR

## DISCUSSION

WGS has been crucial not only to elucidate the genetic relatedness between and within *Arcobacter* species but also to better understand their metabolic pathways, potential pathogenicity and antibiotic resistance determinants [40,65–69]. For *A. butzleri*, which is the *Arcobacter* species causing more concern for human health due to its emergent character [7, 70], WGS has been so far performed for single strain characterization, for the selection of typing loci and for small-scale comparative genomics [40,67,68,71,72]. This has hindered a clear assessment of the evolutionary dynamics and diversification mechanisms of this emerging pathogen. In this context, we performed a deep genome-scale comparative analysis of 49 strains (all publicly available genomes at the study start plus 22 newly-sequenced strains from our bank collection), focused on decoding *A. butzleri* core- and pan-genome diversity and specific genetic traits underlying its pathogenic potential and diverse ecology. We found a scenario of high genomic diversity marked by an open pan-genome overrepresented by strain-specific accessory genome loci and a highly plastic repertoire of virulence- and adaptation-associated determinants.

The hallmark example of how plastic *A. butzleri* can be is the polymorphism pattern of the gene encoding the porin, adhesin and main antigen PorA. In fact, this pathogen is able to exchange *porA* as a whole and/or its antigenic-encoding regions separately, leading to a multitude of chimeric PorA presentations. This diversification pathway can not only markedly impact *A. butzleri* host/environment interaction (e.g. through rapid antigenic shifts), but also potentiate the emergence of variants with enhanced capacity to infect specific hosts, including humans. For instance, in *C. jejuni*, discrete point mutations in a single surface-exposed loop (loop 4, corresponding to the most variable domain of *A. butzleri* PorA) (Fig. 6A), were both necessary and sufficient for causing abortion in the guinea pig model, linking PorA changes to the emergence of a hypervirulent *C. jejuni* clone that is able to cause abortion in ruminants and foodborne disease outbreaks in humans [46]. Again resembling *C. jejuni* [49, 73], in the present study, we identified interstrain in-length variable SSRs potentially leading to ON-OFF switching of protein status or to transcriptomic changes. Concordantly, these SSR affected genes involved in biological mechanisms (LPS modification and chemotaxis) highly linked to the ability of the bacteria to rapidly respond to environmental changes through phase-variation (Table 1). Altogether, these two novel findings reinforce the high adaptive capacity of this ubiquitous bacterium, while also opening new research lines focused on understanding how these genetic mechanisms shape *A. butzleri* virulence traits, transmission capability and ecological success. For instance, it is expected that future large-scale genomic studies will be key to decipher whether individual HVRs profiles (or combined epitope mosaicisms) in PorA or specific status of phase-variable genes correlate with *A. butzleri* population diversity and/or ecology distribution. For example, will the observed incongruence between *porA* and the genome background remain upon the knowledge of the species population structure? Are there specific HVRs patterns linked to human infection? Is there an interplay between specific protein status and specific bacterial niches, environment versus animal host? Besides these novel findings, our screening of potential virulence factors, including both those previously described [40] and newly identified determinants, unveiled not only a virulence arsenal shared by all strains but also several virulence factors found to be unevenly distributed within the species. Again supporting *A. butzleri* hypervariable and plastic character, most strains presented unique virulence repertoires, raising the hypothesis that this human pathogen may harbor an open virulome. With our still limited but diverse dataset, we could consolidate the previous assumption [40,67,68,71,72] that *A. butzleri* is also characterized by the high abundance of genes associated with environment-interacting bacterial functions playing a central role in *A. butzleri* adaptation and survival. In fact, all strains present the complete set of flagellar-related genes, the CheA-CheY chemotaxis system (including the presence of three *cheY* genes) and a multitude of genes encoding methyl-accepting chemotaxis proteins (typically >20 per genome). Besides other previously described virulence determinants here found to be shared by all strains (e.g. *ciaB* and *tlyA*), we highlight the detection of additional common virulence factors based on their homology with *Campylobacter* spp. [42], namely *luxS*, *htrA*, *iamA* and *fur*. All these core virulome genes constitute prioritizing study targets to understand, for instance, how *A. butzleri* promotes cell adherence/invasion and/or subversion of host functions during human infection.

The virulence determinants found to be differentially present among the studied strains revealed a distribution generally not correlating with genome-based phylogeny. This launches the hypothesis that the accessory virulome is hypervariable throughout *A. butzleri* population, with genes circulating between strains regardless of their genetic relatedness and shared ecology. For instance, the urease cluster *ureD(AB)CEFG*, which may be important in the context of host infection, as observed in *H. pylori* [74], was found in 25 strains with rather different genome backbones and isolation sources. We could confirm the urease-positive phenotype for the strains in our collection and potentially link specific mutational patterns in *ureAB* and *ureC* to the lack of phenotype found in one strain carrying the complete urease cluster. Additionally, *hecA* and *hecB* (coding for a filamentous hemagglutinin and a hemolysin activation protein), were found in 13 genomes. These genes are usually PCR-tested to infer *A. butzleri* virulence but they are often not detected simultaneously by PCR screening, with *hecA* being less frequently reported [21,56,58–60]. This may be justified by the *hecA* hypervariable pattern observed in the present study as it yields low primer specificity while challenging *in silico* WGS-based detection by direct nucleotide homology. We anticipate that the upcoming era of genome-based surveillance will certainly demand a revisit and an update of rapid molecular typing tests. For instance, the *in silico* prediction of MLST profiles will also require the development of novel bioinformatics tools to overcome the presence of two non-identical copies of one of the MLST loci (*glyA* loci) [29], a hurdle also observed in other pathogens when transiting to a WGS-based typing [75].

Resistance to multiple antibiotics has been extensively reported in *A. butzleri* [76]. However, this multiresistant character and the underlying mechanisms have been poorly explored to date. Here, we identified a remarkably large set of efflux-pump related genes, with 19 EP systems from different families being identified among the genomes. The fact that ten of these systems are common to all the studied strains raises the question of whether the core of EP-related genes might be contributing to the wide spread resistance of this species to certain antibiotics [76]. Here, by a thorough analysis, a correlation was found between erythromycin resistance phenotype and premature stop codons (SNP-, indel- and IS-mediated) in a putative transcriptional regulator of the TetR Family of Transcriptional Repressors that likely regulates the efflux system EP16 (Fig. 4, Fig. 5). This is, not only, the first description of a genetic determinant of erythromycin resistance in *A. butzleri*, but also the first observation of IS-mediated mechanism of antibiotic resistance in this bacteria. We hypothesize that TetR truncating mutations lead to the overexpression of EP16, to an increase of erythromycin extrusion and ultimately to resistance/tolerance to this antibiotic. This is concordant with the previous description of several efflux pumps as macrolide exporters [77] and contrasts with other resistance mechanisms reported in several bacterial species as *Campylobacter* spp [78], namely mutations in the 23S rRNA, in ribosomal proteins coding genes or the presence of *ermB* (these markers were not found among the screened *A. butzleri* genomes from our collection). In the present study, we also linked fluoroquinolones (ciprofloxacin and levofloxacin) to the presence of the Thr-85-Ile substitution in GyrA, as previously described [79]. Regarding the quinolone, nalidixic acid, all strains were resistant to this antibiotic so, in the absence of a susceptible strain, no association could be established with typical determinants (*gyr* mutations) or other suggested mechanisms, such as hydrophobic quinolones uptake [40]. A strong correlation was also found for ampicillin resistance and the presence of an OXA-15-like β-lactamase, justifying the phenotype of 10 resistant and likely two borderline strains (Fig. 5, Table S1). This association is also supported by the decrease of the resistance rate to amoxicillin when combined with the β-lactamase inhibitor clavulanic acid [76]. Of note, the lack of phenotype in two β-lactamase-harboring strains is intriguing and requires further investigation.

Remarkably, we found one strain, D4963, harbouring a novel T4SS displaying a typical architecture of VirB/D4 secretion systems, which markedly differs from the previously described plasmid-encoded T4SS in *A. butzleri* [80]. T4SSs are well-known virulence mediators in several important human pathogens (e.g. *H. pylori*, *Legionella pneumophila*, *Brucella* spp. or *Bartonella* spp.) namely due to their role in cell-to-cell interaction through the secretion of toxic effector proteins involved in the subversion of key host cell functions [81, 82]. For instance, VirB/D4 T4SSs have been found to be involved in modulation of apoptosis and promotion of interbacterial competition in other bacteria [81,83–85], making this novel finding in *A. butzleri* of particular interest for further investigation.

Another interesting observation was the potential correlation between the presence of a T1SS-like cluster (putatively involved in agglutinin secretion) and an extended repertoire of EPs, which was mostly observed for strains displaying phylogenetic clustering. The T1SS-harboring strains present at least 17 out of the 19 Eps identified, when compared with the detection of 11 to 17 (mean of 14.7) EPs in T1SS-negative strains, with most EP-enriched T1SS-positive strains being also urease-negative, possessing *hecAB* and additional resistance determinants (Fig. 3, Fig. 4). Despite this discrete association with phylogeny, the observed overall lack of correlation between the genome phylogeny and specific virulome/resistome signatures was not unexpected at this dataset level, considering the observed high genomic plasticity and diversity of *A. butzleri.* Nonetheless, the multiple strain- (e.g. the novel T4SS, some EPs and capsule related genes) and group-specific traits (e.g., the EP-enriched T1SS-positive signature, urease cluster, *hecAB*, PorA mosaics) detected in this study will certainly deserve a deep investigation (e.g. to explore its potential association with pathogenicity, specific ecological niches, etc.) when a more complete picture of the global *A. butzleri* population structure and molecular epidemiology could be achieved by large-scale genomic studies.

In summary, the present study constitutes a turning point on *A. butzleri* comparative genomics field. Indeed, it provides extended data on core and accessory genome dynamics and a detailed gene-by-gene functional assignment and allelic profiling, while revealing novel findings supporting that the intraspecies virulome and resistome diversity, the antigenicity modulation and puzzling structure of the main antigen PorA and potential phase-variation mechanisms are key to shape *A. butzleri* environmental/host adaptation and pathogenicity. In another perspective, our results largely reinforce the pathogenic potential of *A. butzleri*, highlighting the need to implement more robust species-oriented diagnostics and molecular typing strategies towards an enhanced global surveillance and control of this emerging human gastrointestinal pathogen.

## AUTHOR CONTRIBUTION

SF collected, cultured and phenotypically characterized the strains. JI did NGS procedures and bioinformatics analyses. MP and VB supported the bioinformatics analyses. FD, MO and JPG contributed to study design and provided all resources for study development. SF, MO, JI and VB wrote the Article. All authors analysed and interpreted the data, and revised and approved the Article. VB and SF jointly supervised the study.

## CONFLICTS OF INTERESET

The authors declare that they have no competing interests.

## ETHICAL STATEMENT

No human experimentation is reported.

## ACKNOWLEDGMENTS

We thank Armélle Ménard from University of Bordeaux for kindly provided the clinical strain Ab_1426_2003.

## DATA BIBLIOGRAPHY

1. Isidro J et al, ENA (reads), PRJEB34441 (2019).

## FIGURES AND SUPPLEMENTAL MATERIAL

**Figure S1.**
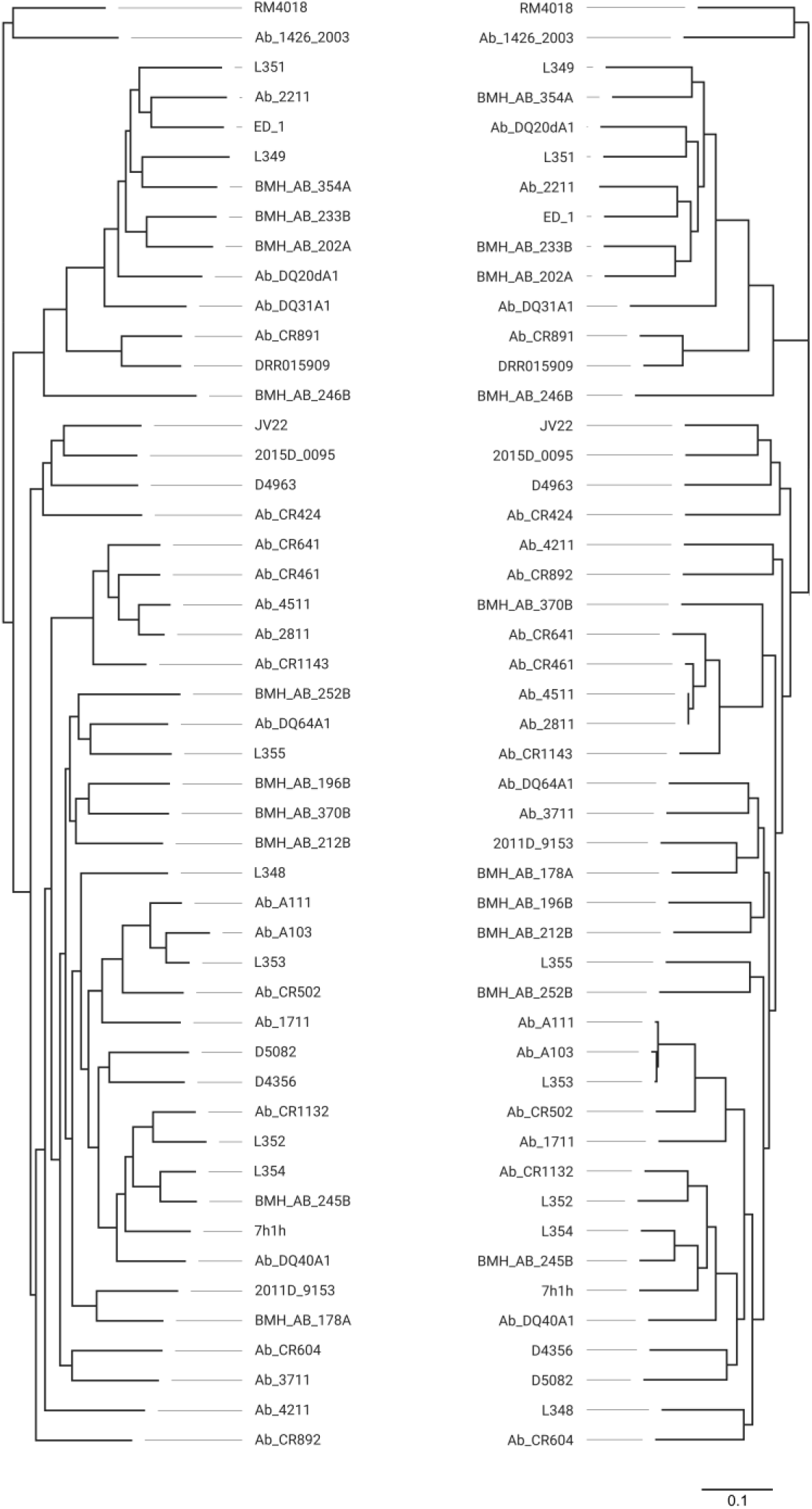
Comparison between the core-genome SNP-based phylogeny of *Arcobacter butzleri* before (left; based on 130195 variant sites) and after (right; based on 114777 variant sites) removal of predicted recombinant regions.

**Figure S2.**
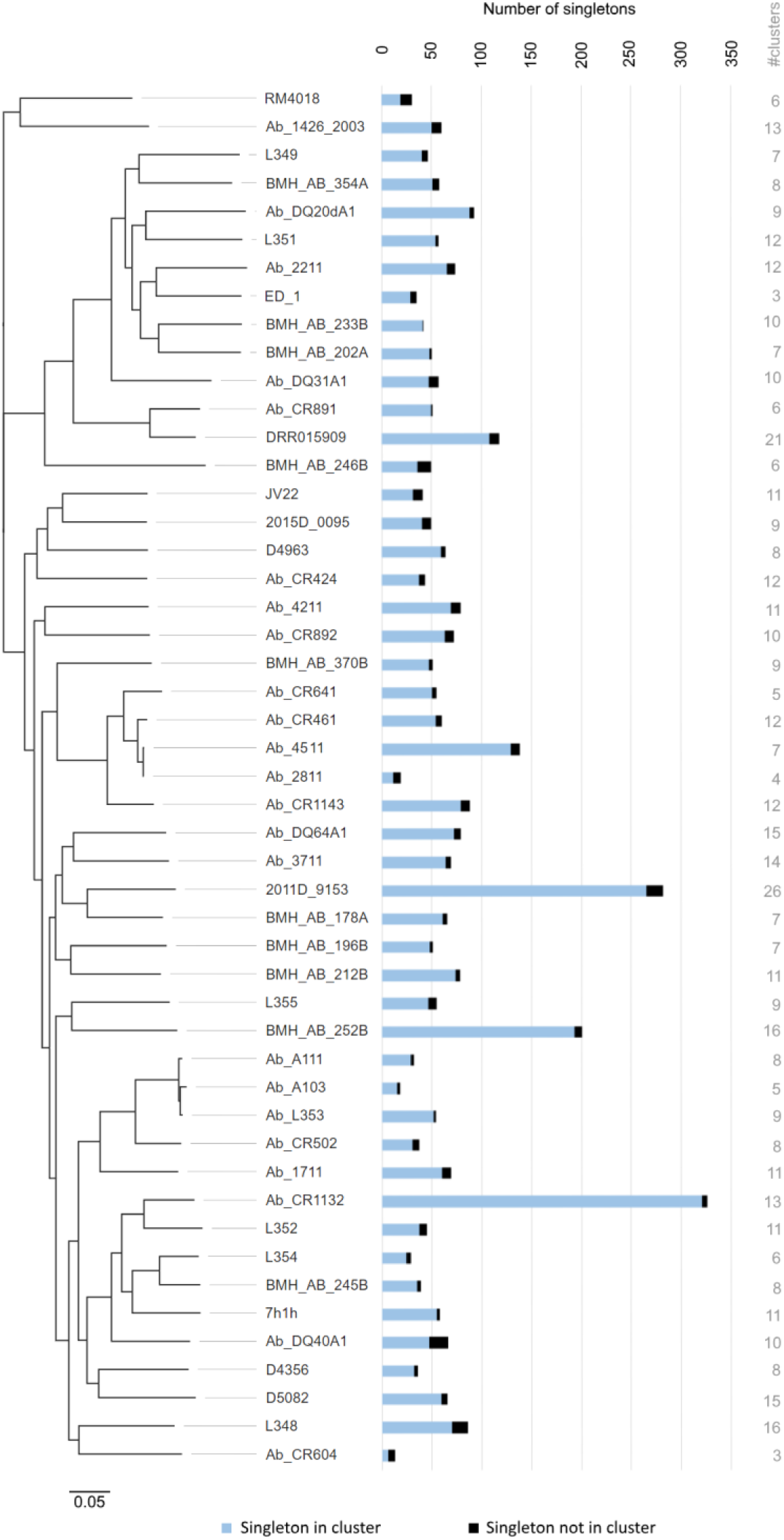
Number and contiguity of the singletons (n=3433) present in each *Arcobacter butzleri* genome. Blue bars represent the number of singletons in cluster (i.e., when at least two singletons are found contiguously in the same genome), while black bars depict the number of singletons not found in cluster. The grey numbers in front of each bar represent the total number of clusters of singletons found in each genome (see methods for details). The functional assignment of all singletons is depicted in Fig. 2B.

**Figure S3.**
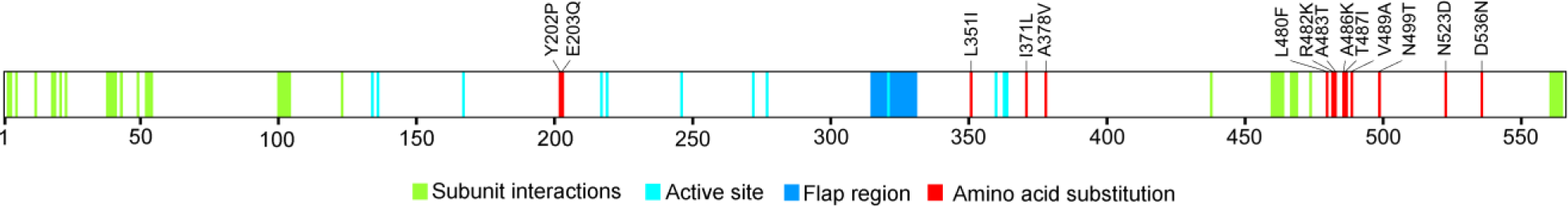
Illustration of the *ureC*-encoded urease alpha subunit of the strain Ab_CR891. The predicted amino acid substitutions highlighted in red were exclusively found in the Ab_CR891 genome [among the *ureD(AB)CEFG*-positive strains] and may justify the negative phenotype obtained for this strain in the urease test. The conserved residues (highlighted in green and blue) are shown as determined by the NCBI Conserved Domain database.

**Figure S4.**
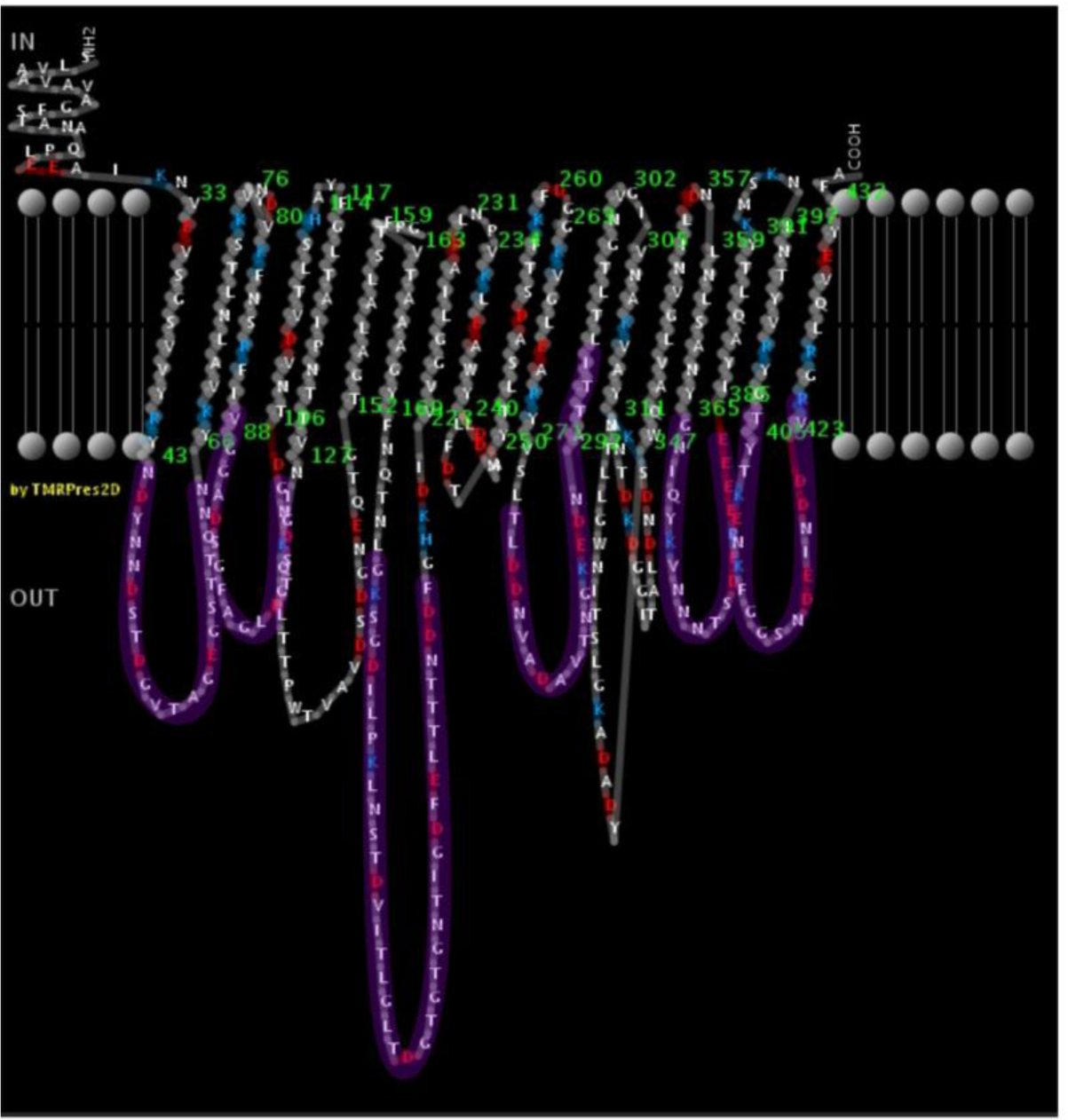
Predicted secondary structure of PorA (using the PorA of *Arcobacter butzleri* reference strain RM4018) and localization of the six hypervariable regions analyzed (highlighted in purple). See methods for details.

**Figure S5.**
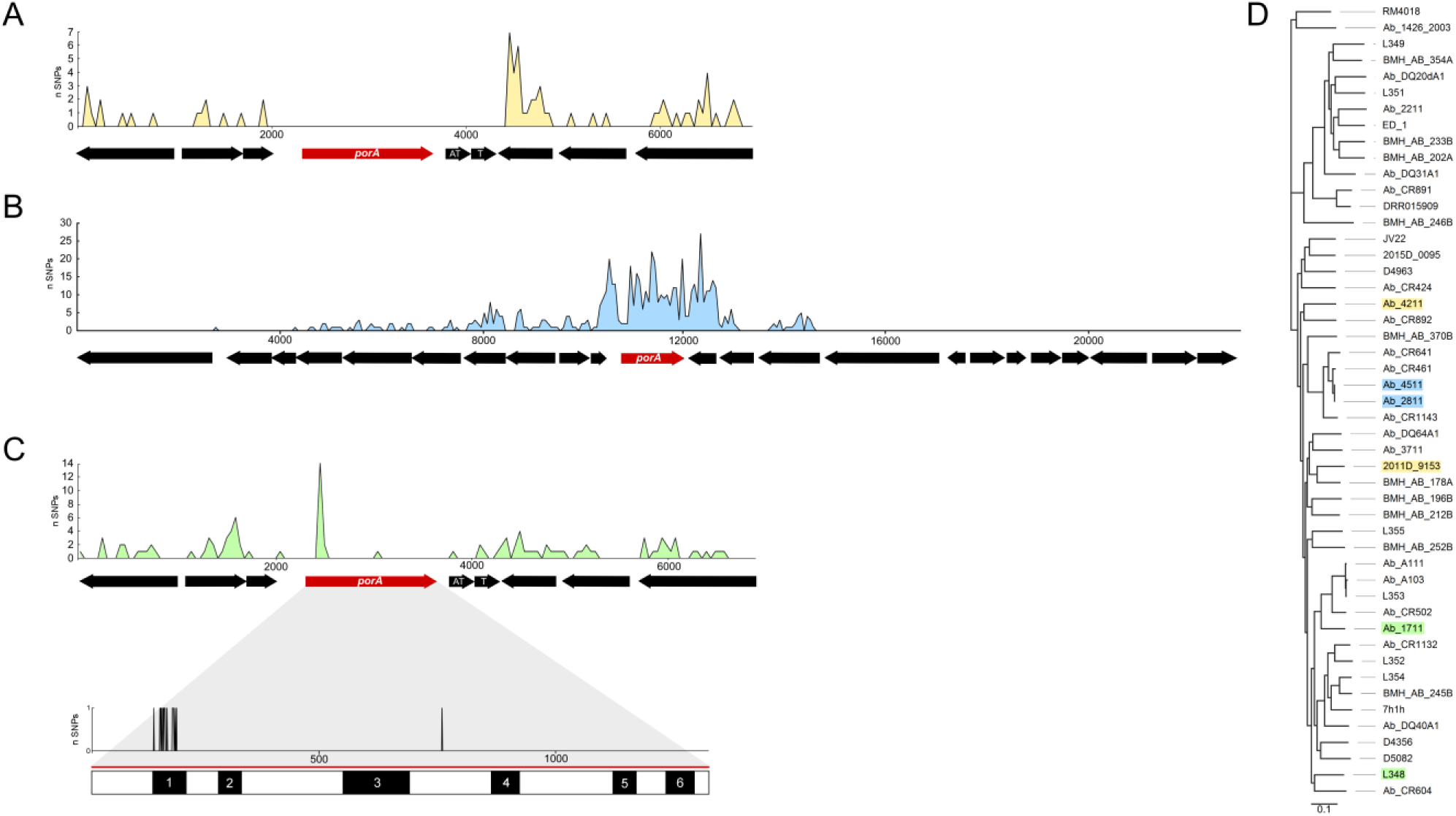
Evidence of the exchange of PorA as a whole and/or its hypervariable antigenic-encoding regions individually. (A) Example of two phylogenetically distant strains, Ab_4211 and 2011D_9153, sharing the same *porA* sequence along with two adjacent genes coding for a toxin/anti-toxin (T/AT) system, while presenting very distinct flanking regions. (B) Example of two phylogenetically close strains, Ab_4511 and Ab_2811, markedly differing in their *porA* (and adjacent regions), while presenting flanking regions with no polymorphism. (C) Example of two phylogenetically distant strains, Ab_1711 and L348, sharing highly similar *porA* sequences that, however, differ markedly in the first hypervariable region (HVR1). The DNA polymorphism analysis was performed with DnaSP v6.10.01 and the single nucleotide polymorphisms (SNPs) are shown with window lengths and step sizes of 45-bp for (A) and (C, larger region), 60-bp for (B), and 1-bp for (C, only *porA*). (D) The six genomes being compared are highlighted in the core-genome SNP-based phylogeny (based on 114777 variant sites detected after recombination removal), according with the colors used in the DNA polymorphism plots (A), (B) and (C).

**Table S1.** *(see Excel file)*

**Table S2.** *(see Excel file)*

**Table S3.** *(see Excel file)*

**Table S4.** *(see Excel file)*

